# Generation of a microsporidia species attribute database and analysis of the extensive ecological and phenotypic diversity of microsporidia

**DOI:** 10.1101/2021.02.21.432160

**Authors:** Brandon M. Murareanu, Ronesh Sukhdeo, Rui Qu, Jason Jiang, Aaron W. Reinke

**Author notes:** These authors contributed equally (Brandon M. Murareanu, Ronesh Sukhdeo, Rui Qu, Jason Jiang).

## Abstract

Microsporidia are a large group of fungal-related obligate intracellular parasites. Though many microsporidia species have been identified over the past 160 years, there is a lacking depiction of the full diversity of this phylum. To systematically describe the characteristics of these parasites, we created a database of 1,440 species and their attributes, including the hosts they infect and spore characteristics. We find that microsporidia have been reported to infect 16 metazoan and four protozoan phyla, with smaller phyla being underrepresented. Most species are only reported to infect a single host, but those that are generalists are also more likely to infect a broader set of host tissues. Strikingly, polar tubes are 3-fold longer in species that infect tissues besides the intestine, suggesting that polar tube length is a determinant of tissue specificity. Phylogenetic analysis revealed four clades which each contain microsporidia infecting hosts from all major habitats. Although related species are more likely to infect similar hosts, we observe examples of changes in host specificity and convergent evolution. Taken together, our results show that microsporidia display vast diversity in their morphology and the hosts they infect, illustrating the flexibility of these parasites to evolve new traits.

## Introduction

Microsporidia are a large phylum of obligate intracellular parasites. Although they were once thought to be protists, it is now clear they belong to a lineage of early diverging fungi(1). Microsporidia was the first parasite to have its genome sequenced, revealing the smallest eukaryotic genome with as few as ~2000 protein coding genes(2, 3). Genome reduction results from the loss of many metabolic and regulatory genes, reduced non-coding DNA, and reduced gene sizes that encode for proteins smaller than their yeast orthologs(4–7). Microsporidia have also lost mitochondria and instead depend on reduced organelles known as mitosomes(8). As a result, microsporidia rely heavily on their hosts for nutrients and have evolved many novel proteins for interacting with host cells(9, 10). Paradoxically, although microsporidia are amongst the simplest eukaryotes, there are at least 1500 known species that as a phylum can infect many different animals and most major tissues(11, 12).

Microsporidia cause death and disease and are a threat to both animal and human health. They were first discovered in 1857 in silkworms and microsporidia infections have triggered historical collapses of the sericulture industry(13). Microsporidia infections in animals are very common. For example, 50% of honey bee colonies and 60% of field-collected mosquitos have been reported to be infected with microsporidia(14, 15). These parasites are an emerging threat, with the most common species that infect honey bee colonies and shrimp farms only being discovered within the last two decades(16, 17). Microsporidia also cause problems in laboratory research, commonly infecting zebrafish facilities and causing fruit fly colonies to collapse(18, 19). Infections often result in delayed animal growth, reduction in offspring, and premature mortality(20–23). Over a dozen species of microsporidia can infect humans and several of these species are also found in a large variety of different mammals and birds, leading to concern over zoonotic transfer(24–26). In addition to the threats posed by microsporidia, they also have potential benefits, including being used as biocontrol agents against insect pests or preventing the spread of mosquito-vectored diseases(27, 28).

All microsporidia produce environmentally resistant spores. These spores contain both the infectious sporoplasm and the unique polar tube infectious apparatus(29). In the presence of a host, microsporidia fire their polar tube which is thought to either pierce the host cell or fuse with the host plasma membrane to form an invasion synapse(30). Each species has spores of a characteristic size, but some species have multiple classes of spores(31, 32). While many infections are thought to result from the oral consumption of spores, other types of infections e.g. ocular, can also occur(26).

Although many microsporidia species have been characterized, their properties have not been examined systematically. Thus, the true diversity of these medically and agriculturally important parasites is unknown. Here, we curate a database of microsporidia attributes for 1440 species. Using this data, we describe the history and geography of microsporidia discovery. We show that microsporidia can infect a diverse range of host species from all major habitats. We find that microsporidia that infect more hosts tend to infect more tissues. We observe large ranges for spore size, which correlates with the number of nuclei and polar tube length. We also find that the length of the polar tube is longer in species that infect other tissues besides the intestine. Phylogenetic analysis shows that microsporidia group into clades with predominant environmental habitats, but the ability to infect in other environments within a clade is also common. Related species mostly infect similar hosts, but dramatic changes in host specificity and convergent evolution also occur. Together, this work describes the diverse properties of microsporidia species and provides a resource for further understanding these enigmatic parasites.

## Results

### Generating a database of microsporidia species’ characteristics

To determine the reported diversity of microsporidia species, we comprehensively examined the literature. We began with Victor Sprague’s 1977 book, the most complete compendium of microsporidia published thus far, which contains information on 692 species(33). We then identified 704 journal articles containing reports of identified microsporidia species. We recorded the described attributes of each species including the date and location of discovery, host species, infected tissues, spore size, and polar tube length. Overall, we generated a database containing information on 1,440 microsporidia species (Table S1).

We first examined the history of microsporidia species discovery. The original article describing the first species of microsporidia was published in 1857(34). Since 1881, on average ~100 novel species have been reported per decade. The number of discovered species peaked in the 1970s with 246 species being reported. Although in recent years fewer species have been reported than at this peak, during the last decade, on average 14 novel species have been described every year (Fig. 1A).

**Fig. 1.**
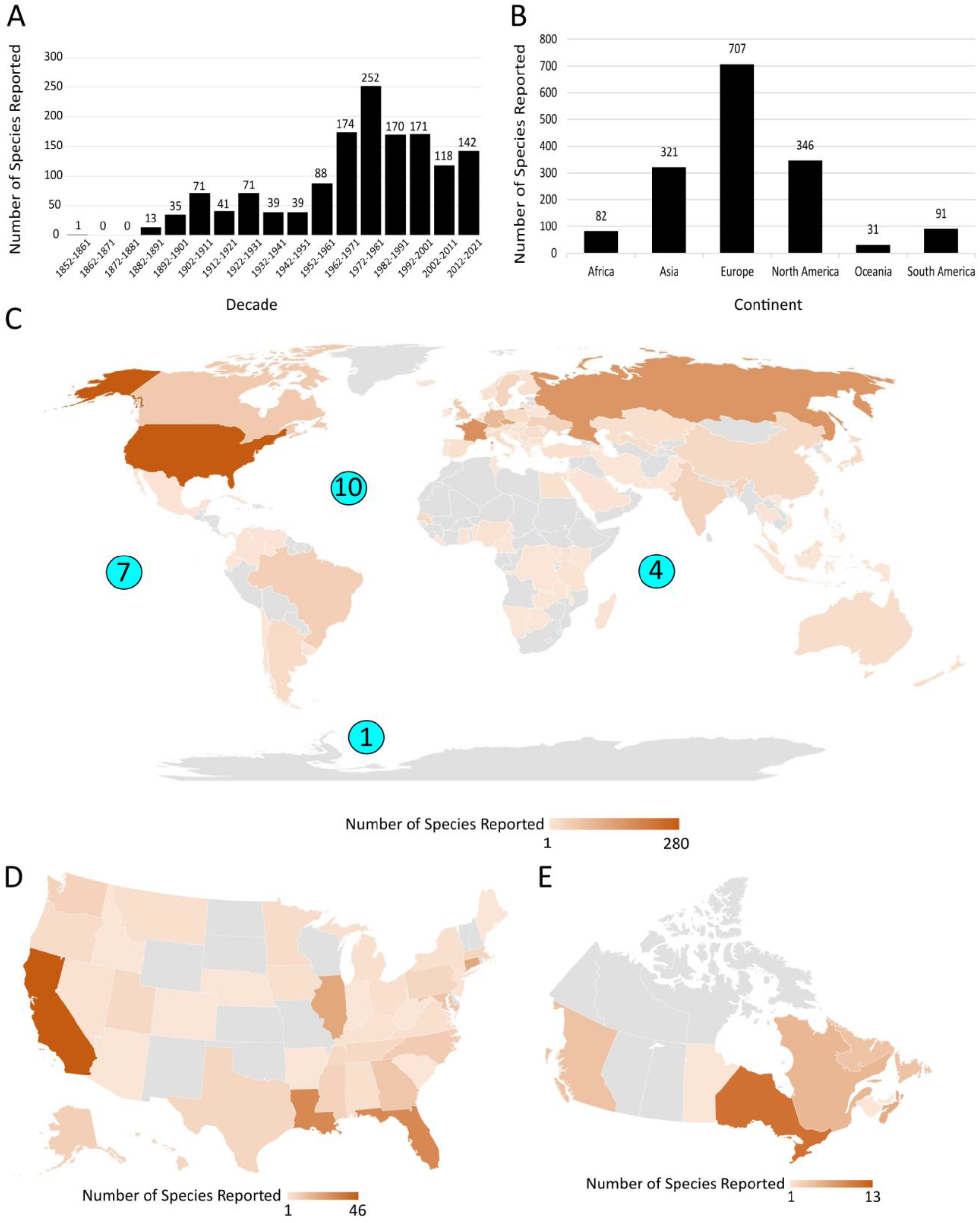
History and geography of microsporidia species discovery. **(A)** Number of new microsporidia species described per decade (n = 1425 microsporidia species). **(B)** Number of microsporidia species found on each continent (n = 1310 microsporidia species). **(C-E)** Geographical location of microsporidia, with each species depicted once for each locality where they have been reported. The number of species found in each locality is depicted as a heatmap, according to the scale at the bottom. Areas in grey indicate that no species have been reported. **(C)** Location of microsporidia species identified, by country and ocean. Numbers in blue circles describe species found in respective oceans (n = 1310 microsporidia species). **(D)** Location of microsporidia species reported by state within the United States (n = 249 microsporidia species). **(E)** Location of microsporidia species reported by province and territory within Canada (n = 45 microsporidia species).

We next determined the geographical locations from which microsporidia have been isolated. Although the majority of described species have been found in Europe and North America, microsporidia have been found in all continents except Antarctica (Fig. 1B). They have also been reported in 11 oceans or seas, including a species found in the Weddell sea off the coast of Antarctica, and from methane seeps on the floor of the pacific ocean(35, 36). While the United States has reported the most species of microsporidia (283), 123 species have also been found in the much smaller Czech Republic (Fig. 1C). We also see a distribution of reported species within a country, with microsporidia being reported in the majority of states and provinces in the United States and Canada (Fig. 1D and E).

### Hosts and tissues reported to be infected by microsporidia

Microsporidia have often been described as infecting nearly all animal taxa, but a full description of which host phyla these parasites infect is lacking(26, 37, 38). Using several databases describing animal and protozoan diversity, we determined the taxonomical placement of 1436 microsporidia hosts (Table S2). We find that infections have been reported in 16 metazoan and four protozoan phyla. The majority of the hosts in six of these phyla are themselves parasites, demonstrating that hyperparasitism is a widespread microsporidian trait (Fig. 2A). There are 16 metazoan phyla that have not been reported to be infected by microsporidia. These phyla have on average 26-fold fewer known species than phyla that have been reported to be infected with microsporidia. This suggests that the absence of known microsporidia infections in these phyla may be due to a lack of sampling from these hosts (Fig. 2B). We also observed a diversity of hosts within a phylum; the two phyla with the most hosts, Chordates and Arthropods, contain hosts in 39 out of 155 orders and 42 out of 162 orders, respectively (Fig. 2C and S1).

**Fig 2.**
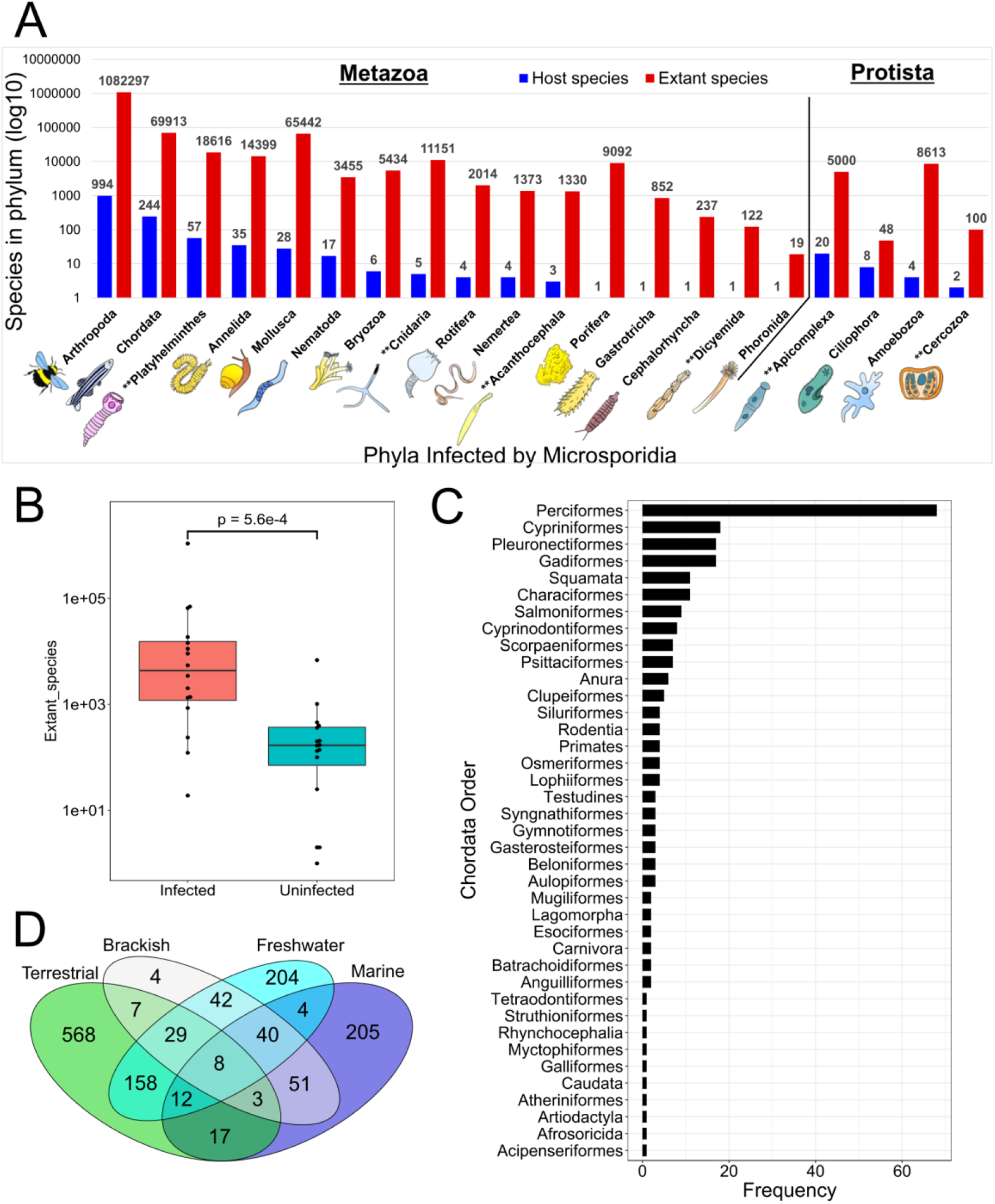
Microsporidia infect a diverse range of hosts from all major environments. Host taxonomy and environment was determined for each reported microsporidia species (see methods). **(A)** Distribution of microsporidia hosts by metazoan (left) and protist (right) phyla. The number of host species in each phylum (blue) and the total number of extant species in each phylum (red) are shown. Phyla where the majority of microsporidia hosts are parasites are indicated (**)(n = 1436 host species). **(B)** Number of described species from phyla reported to be infected by microsporidia (red) (n = 16 phyla) and those that have not been reported to be infected by microsporidia (blue) (n = 16 phyla). The Mann-Whitney U test was used to compare mean ranks between the groups. **(C)** Frequency of Chordate orders reported to be infected by microsporidia (n = 39 orders). **(D)** Venn diagram of the major environments that microsporidia hosts dwell in (n = 1352 host species).

Because microsporidia are obligate intracellular parasites, the habitat of the host informs that of the microsporidia species(12). We determined the ecological environments of the different hosts in our database. Of the four major habitat types, 59.3% of hosts are terrestrial, 36.8% of hosts are freshwater, 25.1% of hosts are marine, and 13.6% of hosts are brackish (of hosts with environment data). Interestingly, 27.4% of hosts with environment assignments are classified as living in more than one environment. This is exemplified by eight hosts, including the mosquito (*Anopheles pseudopunctipennis)* and parasitic flatworm (*Heterophyes heterophyes*), which inhabit all four environment types (Fig. 2D and Table S2).

Microsporidia are known to infect a range of different tissues within their hosts(12). To investigate the extent of this diversity, we classified infections into 23 tissue types (Table S3A). The tissues infected by microsporidia encompass all embryonic germ layers and most major organ systems(39). The adipose tissue, intestine, muscles, germline, and excretory system are the most commonly infected tissues. Infections have also been reported in specialized tissues such as the gills of fish and oenocytes, which are lipid-processing cells in insects(40) (Fig. 3A). Most microsporidia infections appear to be restricted to a single tissue, but ~11% of species can infect four or more tissues (Fig. 3B).

**Fig. 3.**
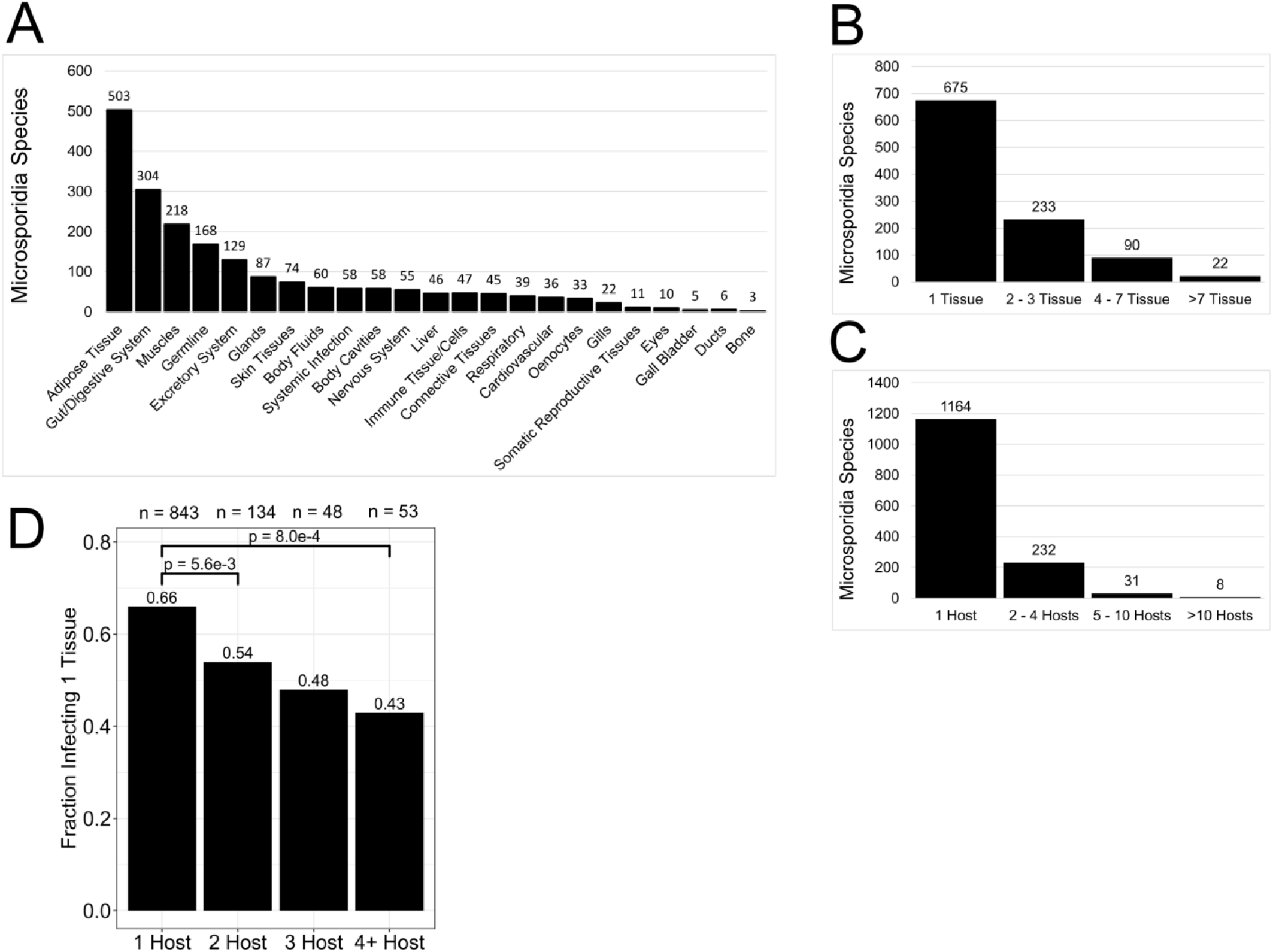
Microsporidia infect a diversity of tissues and species that infect more hosts tend to infect a broader set of tissues. **(A-B)** The tissues described to be infected by microsporidia were placed into 23 tissue-types (see methods). **(A)** The number of microsporidia species reported to infect each tissue type (n = 1123 species). **(B)** Distribution of the number of tissue types infected by each microsporidia species (n = 1020 species). **(C)** Distribution of the number of hosts infected by each microsporidia species (n = 1435 species). **(D)** Proportion of species that infect more than one tissue compared to how many hosts are infected by that species (n = 1078). Species that are described to have general or systemic infections are considered to infect more than one tissue. Significant difference between proportions were determined with pairwise chi-square tests, with Bonferroni p-value correction (α = 0.05/6 comparisons = 8.3 × 10^−3^).

Microsporidia can be specialists that infect only one or a few hosts, or generalists that can infect many hosts(5, 37). Over 80% of reported species (with host data) only infect a single host, but 2.2% can infect five or more hosts (Fig. 3C). To determine if there is a relationship between host range and tissue specificity, we examined if species that infect more hosts also infect more tissues. Indeed, 34% of species that infect one host infect multiple tissues, compared to 57% of species that infect four or more hosts infecting multiple tissues (Fig. 3D). This correlation suggests that species with larger host ranges tend to infect a broader set of tissues.

### Diversity of microsporidia spore morphology

All microsporidia have an extracellular spore form and the morphological properties of these spores can be used to distinguish species(31). To examine the diversity of microsporidia spore morphology, we first classified spores into 20 unique shape descriptors (Table S3B). Over 93% of classified spores belong to just six shape classes (Fig. 4A). Examination of spore size revealed a large diversity in both spore length (1 – 25 μm) and width (0.15 – 10 μm). Although there is overlap between shape description and spore dimensions, there are differences in the average length/width ratio of the main shape classes with rod-shaped being 3.7, ovocylindrical being 2.7, oval, pyriform, and elliptical being 1.8-1.9, and spherical being 1 (Fig. 4B and S2).

**Fig. 4.**
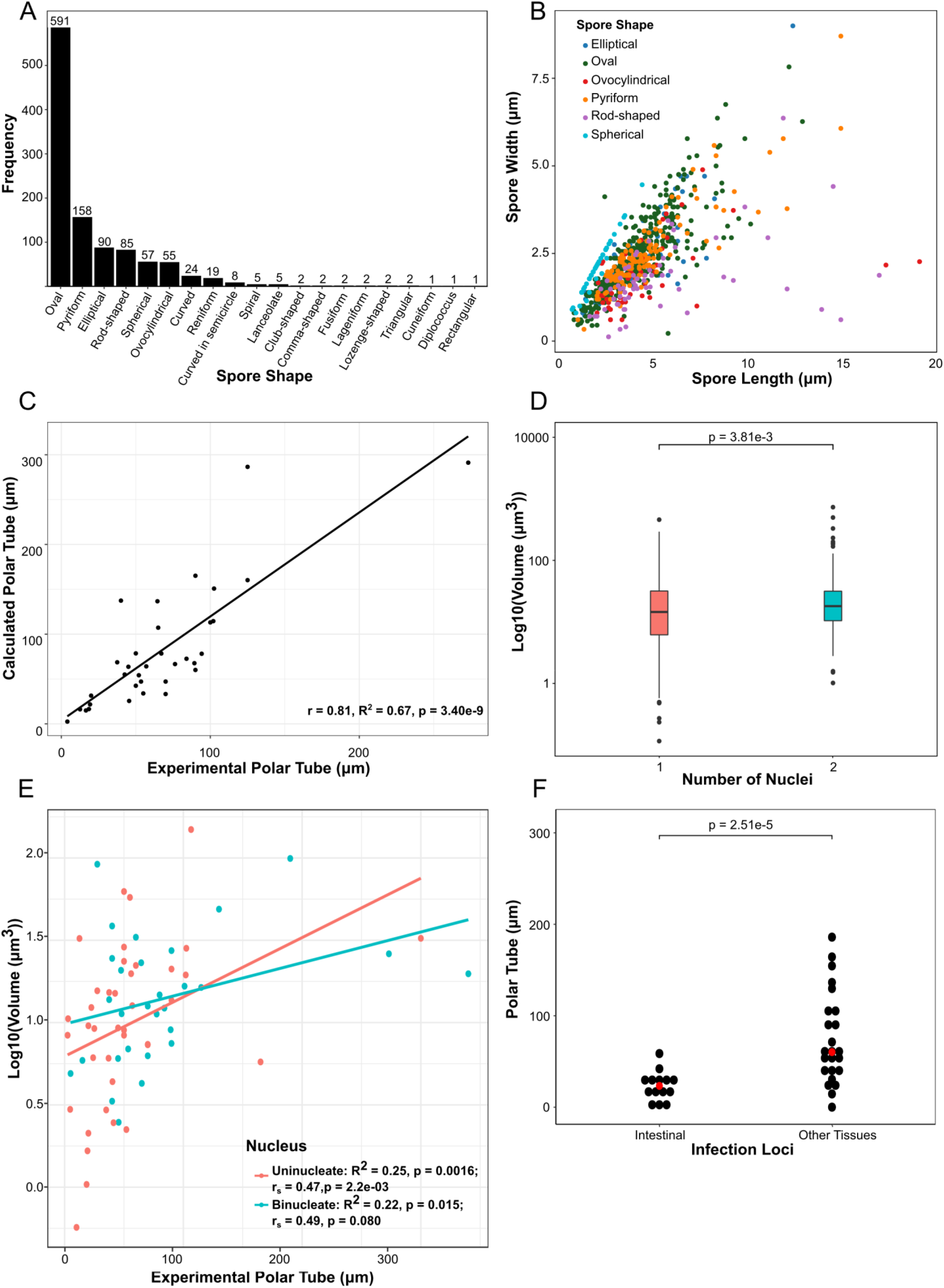
Microsporidia spores display a diversity of shapes, sizes, and polar tube lengths. **(A-B)** Spore shapes were placed into 20 morphology classes (see methods and Table S3B). **(A)** The frequency of each type of spore shape type (n = 1112 spores). **(B)** Spore length and width dimensions plotted for the six most common spore shapes (n = 816 spores). **(C)** Comparison of experimentally determined and calculated polar tubule lengths (n = 34 spores). **(D)** Median spore volume for uninucleate (n = 496) and binucleate (n = 287) spores. The p-value was determined by the Mann–Whitney U test. **(E)** Comparison between experimental polar filament lengths and spore volume (n = 37 spores for uninucleate and 26 species for binucleate). Correlation was calculated with spearman r_s_. **(F)** Relationship between polar tubule length and tissues infected. Median polar tube values are indicated in red. Significance was calculated using a paired T-test between “Intestinal” species and “Other Tissues” species infecting the same host.

Using length and width measurements of spores, we estimated spore volumes, which range from 3.5 × 10^−2^ μm^3^ − 1.3 × 10^3^ μm^3^. We compared our volume estimates with 3-dimentionsal measurements using the two species for which such data is available(41). For *Anncaliia algerae*, we estimated a volume of 22.7 μm^3^ and the volume was calculated to be 8.8 μm^3^. For *Encephalitozoon hellem*, we estimated a volume of 2.1 μm^3^, and the volume was calculated to be 3.6 μm^3^. We also used the same volume formula, but the length and width measurements from (41), and estimated *A. algerae* to be 12.8 μm^3^ and *E. hellem* to be 4.1 μm^3^. These comparisons demonstrate, at least for this limited set, that our estimated volumes are similar to those calculated using three-dimensional methods and that the majority of the differences in volume arise from differences in spore length and width measurements.

Microsporidia spores contain polar tubes that fire in the presence of a host signal and are used to infect host cells(30). The length of the polar tube can be measured by artificially triggering spore firing by treating with different chemicals or mechanical pressure(30, 42). The polar tube is coiled inside of the spore and the number of coils can be determined using electron microscopy(29). Using the spore diameter and the number of polar tube coils, we estimated the length of the unfired polar tube. We observe a large range in both experimentally measured (2.5 μm – 420 μm) and estimated (1.5 μm – 752.1 μm) polar tube lengths. We compared estimated polar tube length to calculated polar tube length and observed a significant correlation (Fig. 4C).

Microsporidia spores need to be able to accommodate both the polar tube and the sporoplasm. Sporoplasm size measurements do not exist for most species, so we used the number of nuclei (either one or two) as a proxy for the size of the sporoplasm. We examined the relationship between spore size and nuclei number and observed that spores with two nuclei are on average ~22% larger than those with one nucleus (Fig. 4D). We next examined the relationship between polar tube length and spore size. We observed that experimentally measured polar tube lengths were positively and significantly corelated with spore volume (Fig. 4E and S3). Taken together, these results suggest that although the number of nuclei and the length of the polar tube do not explain all of the variation in spore volume, the size of intracellular contents is correlated with spore size.

One possible explanation for why polar tube length varies so widely, is that the polar tube specifies which tissues are infected. This idea was proposed by Luallen and colleagues studying microsporidia infection in the nematode *Caenorhabditis elegans*(6). The authors observed that *Nematocida parisii*, which infects the intestine, has a polar tube length of 4.03 μm, in contrast to that of *Nematocida displodere*, which infects the muscle and epidermis from the intestinal lumen, and has a polar tube length of 12.55 μm. To determine if polar tube length correlates with infected tissue type, we examined 11 hosts that were infected by at least one microsporidia species that exclusively infects the digestive system and at least one species that infects other tissues. In 82% of microsporidia species pairs, the polar tube was longer in the microsporidia species that infected other tissues besides the digestive system. On average, polar tubes of species that infect other tissues were 3-fold longer than those of species restricted to the digestive system. This suggests that one function of polar tube length is determining host-tissue specificity (Fig. 4F and Table S4).

Microsporidia can either be transmitted horizontality, from host to host, or vertically, from parent to offspring(20, 43). We examined whether species were reported to infect the germline (vertical) or somatic tissue (horizontal) and observed that ~18% of species are potentially vertically transmitted (Fig. S4A). We also observed that transmission mode is not dependent on host environment (Fig. S4B). Furthermore, spore volume and polar tube length are not correlated with transmission mode (Fig. S4C-D).

### Evolutionary properties of microsporidia

Microsporidia have previously been reported to group into five clades, classified based on host environmental habitat(44). Using our large collection of host and species information, we sought to revisit these groupings. We performed Bayesian phylogenetic analysis on 273 18S rRNA sequences belonging to microsporidia species for which we also had attribute data (Table S5A and B). The resulting tree recovers four clades (I,III,IV, and V) and is largely in agreement with recently published microsporidia phylogenetic trees(45, 46) (Fig. 5A and S5). We also assembled a Maximum Likelihood phylogenetic tree using the same sequences, and the majority of species (~95%) were placed in the same clade in both trees, further supporting our clade groupings (see methods). All subsequent analysis was done using the Bayesian tree. We then determined the percentage of microsporidia species in each clade that inhabit each of the four major environments. We mostly observed the previously determined environmental groupings, with clade IV being terrestrial and clade III being marine. We also observed that clades I and V have a higher percentage of microsporidia with freshwater hosts than the other clades, as has been previously shown. However, clades I and V have a higher percentage of microsporidia with terrestrial than freshwater hosts (Fig. 5B). Although there is a predominant habitat in each clade, between 29-44% of microsporidia species in each clade do not infect a host in the predominant habitat. This suggests that evolution of microsporidia within clades is not strictly restricted to hosts from a single environmental habitat.

**Fig. 5.**
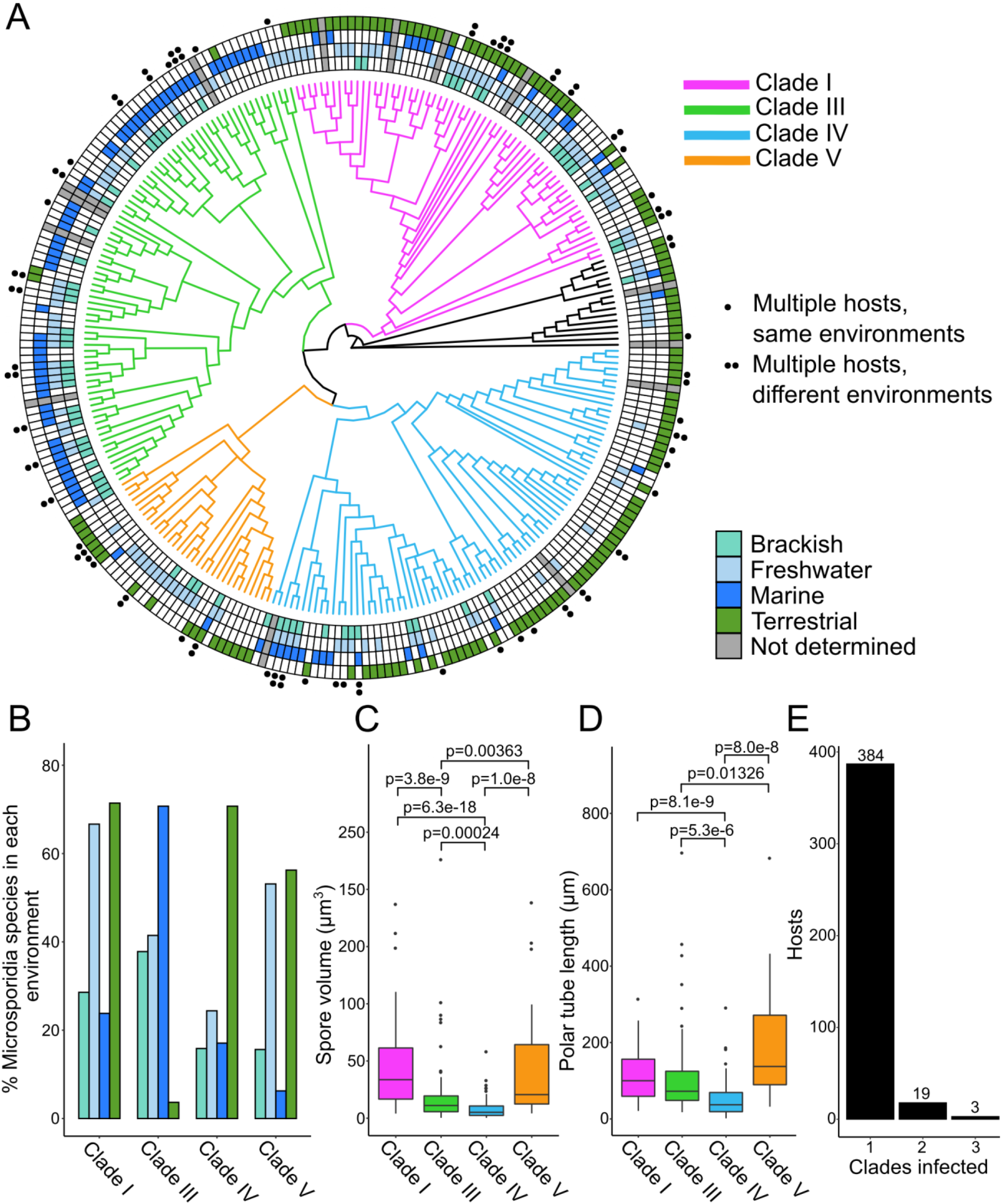
Bayesian 18S Phylogeny of Microsporidia Species Reveals Four Phenotypically Diverse Clades. **(A)** 18S phylogeny of 273 microsporidia species, inferred under a Bayesian framework (see methods). Clades are colored and number of hosts indicated according to the legends at the right. Species not belonging to any of the four main clades are colored black. **(B-D)** Analysis of microsporidia phenotypes in each of the four major clades. **(B)** Frequency of host environments by clade. **(C)** Spore volume distribution by clade. **(D)** Calculated polar tube length distribution by clade. Significance evaluated using pairwise Mann-Whitney U-tests, with Bonferroni p-value adjustments applied to correct for multiple testing (α = 0.05/6 = 8.3 × 10^−3^) **(E)** Number of hosts infected by microsporidia species from one, two or three different clades (n = 406 microsporidia hosts).

We next examined the diversity of spore size and polar tube length amongst clades (Fig. 5C and D). We observe that the clades with the highest percentage of microsporidia infecting freshwater hosts (I and V) have on average larger spores and more variation in spore size than the other two clades (Fig. 5C). This same trend exists in polar tube length, but to a lesser extent (Fig. 5D).

Related species of microsporidia are known to infect very different hosts, but it is unclear how common of an occurrence this is(7). To determine the relationship between microsporidia species similarity and host similarity, we first calculated the pairwise distances between 270 microsporidia 18S rRNA sequences (Fig. 6A). We then compared how often related species infect the same host family, order, class, or phylum. We observed that while 43% of microsporidia species pairs with greater than 65% 18S similarity infect hosts from the same family, this was true for only 2% of pairs with less than 65% 18S similarity. These strong associations between microsporidia and host relatedness were also observed at the other taxonomic ranks (Fig. 6B). Although very closely related microsporidia almost always infect the same phylum, we observed 28 pairs of species with greater than 80% similarity infecting different phyla (Table S5C). This data includes known examples of highly similar species infecting different phyla such as between microsporidia that infect ciliates and those that infect aquatic arthropods and between microsporidia infecting aquatic crustaceans and those infecting parasitic trematodes and paramyxids (47, 48). The species pairs we identified also included eight such instances between species that infect arthropods and the known human infecting species *Vittaforma corneae* and *Trachipleistophora hominis* (26). To identify examples of convergent evolution, we determined how often a host species is infected by microsporidia from more than one clade and found 22 such cases (Fig. 5E). Taken together, our results indicate that microsporidia mostly evolve to infect similar hosts but can undergo dramatic hosts shifts.

**Fig. 6.**
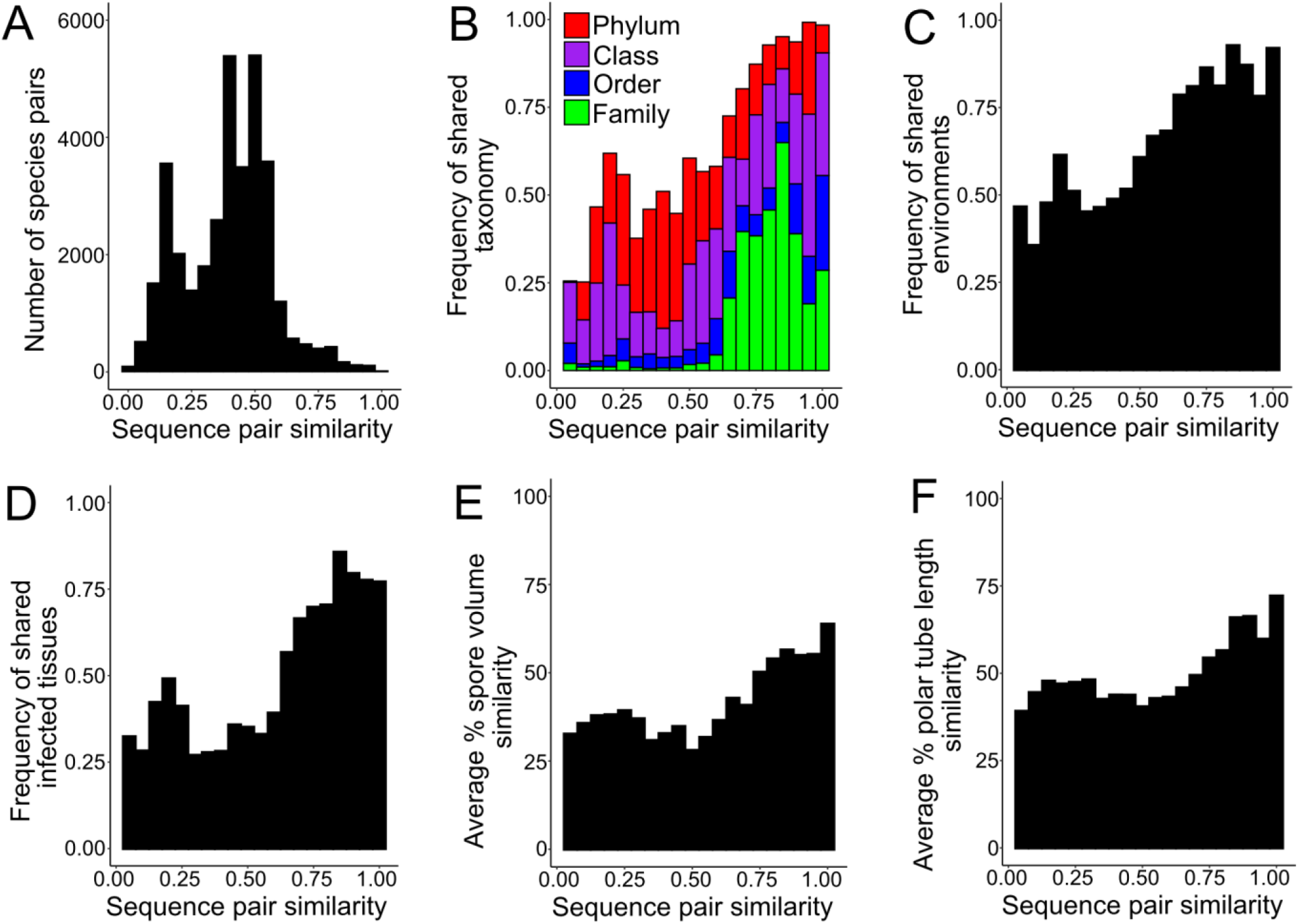
Evolutionary relatedness of microsporidia phenotypes. **(A)** Distribution of species pairs according to sequence similarity (n = 36,315 comparisons). **(B-F)** Distribution of sequence pair similarity and frequency of shared **(B)** host taxonomy (n = 34,980 comparisons), **(C)** environments (n = 33,670 comparisons), **(D)** infected tissues (n = 27,028 comparisons), **(E)** difference in spore volume (n = 31,878 comparisons), **(F)** difference in calculated polar tube length (n = 17,955 comparisons).

Finally, we determined the relationship between microsporidia sequence relatedness and infected tissues, habitat, and spore similarity. We find that closely related species mostly infect the same tissues and share the same environmental habitats (Fig. 6C-D). Spore size and polar tube length are also more similar in closely related species, but to a lesser extent than that of host similarity (Fig. 6E-F). Even closely related species show less than 75% similarity in the tissues they infect and in their spore properties. These results suggest that although related species are likely to have shared characteristics, microsporidia are flexible in their ability to evolve new traits.

## Conclusions

Here we report the largest systemic collection of microsporidia species characteristics assembled since Sprague in 1977(33). Although the data we present is comprehensive, there are several limitations. First, we only curated named species that were reported with attribute data. There are at least 93 identified microsporidia species that we could not find information on and therefore did not include in our analysis(49). Second, many microsporidia species names have changed throughout the years and this likely led to missing information(50). Third, we do not include complete attribute data for every species as authors do not always report on the same properties. Fourth, the data we collected is from a variety of different sources and some characteristics such as spore size were determined through multiple techniques with differing accuracy, lending some uncertainty to the data and our analyses. Finally, for species that are the subject of many publications, we are unlikely to have captured all of the known data including the full breadth of hosts these species infect.

About 1500 microsporidia species have been identified, which is likely only a small fraction of the true number of species(51). We find that identification of novel microsporidia species peaked in the 1970s, which is consistent with a previous report showing identification of microsporidia has lagged behind other parasites(52). Sampling for microsporidia species has been biased with ~65% of the species identified so far being from Europe and North America. Africa, South America, and Oceania have been shown to have large levels of parasite-host diversity, suggesting that these parts of the world could contain many undiscovered microsporidia species(53).

Although the pace of microsporidia discovery has slowed, many new and diverse species are still being discovered. Since the 1990s, microsporidia were reported for the first time in seven phyla (Cephalorhyncha, Porifera, Amoebozoa, Dicyemida, Gastrotricha, Phoronida, and Nemertea), suggesting that most types of animals may be infected by microsporidia. In support of this, although there are no named microsporidia species infecting tardigrades, animals in this phylum has been observed to be infected by microsporidia(54). Environmental sampling has also revealed the existence of many species of unnamed microsporidia, especially a diversity of short-branched microsporidia(45, 55, 56). Echinoderms comprise the largest phylum without any reported microsporidia infections and represent an interesting target for future efforts to uncover novel microsporidia diversity.

The molecular mechanisms that determine host and tissue specificity in microsporidia are mostly unknown. Here we show that microsporidia polar tube length is longer in species that infect tissues other than the intestine. In bilaterian animals, the intestine is in the middle of the animal and access to most other tissues requires going through the intestine(57). This has been shown in *C. elegans*, where infection of the epidermis is dependent upon oral consumption of spores(6). Microsporidia have been shown to be more selective for which types of cells they infect inside of animals than in cell culture (58, 59). Microsporidia have also been shown to infect tissues via injection that were not infected during feeding of spores to mosquito hosts(60, 61). These results suggest that access to tissues could determine specificity and that longer polar tubes could be used to access these other tissues. Microsporidia infections can be disseminated within a host both through spread of meronts between cells and the production of spores that infect other tissues(21, 62). Thus, it is possible that in some cases spores with shorter polar tubes could infect more distal tissues after an initial round of infection. We also show that there is a tendency for microsporidia species that infect more hosts to also infect more tissues. This is likely even more common than our data indicates, as it only takes a single infected animal to determine tissue distribution, but many identifications to determine host range(31). Our data suggest that the evolutionary forces that drive species to be generalists also tends to generate species with broad tissue specificity.

In 2005 Vossbrinck and Debrunner-Vossbrinck proposed that microsporidia compose five major clades that group according to the environmental habitat of each species host(44). Although these environmental groupings provide an attractive explanation of microsporidia evolution, none of the clades only contained hosts from a single environment(63). Clade II was not well supported in this original phylogeny and has not been consistently recovered in more recent phylogenetic trees(20, 31, 45, 55). The position of *Nematocida* in these phylogenies has not been constant and often differs from the basal position recovered from trees based on whole genome sequencing(46, 64). Environmental sampling from diverse environments recovered microsporidia from many different clades, demonstrating that there is not a strict correspondence between clades and host environment. For example, many samples from marine environments were found in clade IV (Terresporidia). Intriguingly, several OTUs (operational taxonomy unit) were found in multiple environments, providing support for some species having multiple hosts from different environments or the same host being infected in multiple environments(45). Our analysis suggests that although clades do belong predominantly to a single environment, microsporidia can evolve to infect hosts in other habitats. A host often exists in more than one habitat and some microsporidia have multiple hosts that exist in distinct habitats. Because of this, many species of microsporidia are found in multiple environments, likely aiding their ability to evolve to infect new hosts.

In conclusion, we have assembled a collection of microsporidia attributes for 1,440 species. This data will be a useful comparative resource which will support further species identification. In addition to the questions that we have examined, this database will aid in addressing other systematic inquires about microsporidia diversity. The community of microsporidia researchers can add to the microsporidia species attributes database by submitting newly discovered species and updating existing species at https://www.reinkelab.org/microsporidiaspecies.

## Materials and methods

### Construction of microsporidia species properties database

Each microsporidia species described from Victor Sprague’s 1977 book was manually entered into an Excel table(33). Each row represented a microsporidium, and columns were created for each species attribute recorded. Parenthesized text was used to denote which spore class or host an attribute corresponded to. The spore conditions (ex: fixed, fresh, etc.) under which morphological attributes were recorded were also denoted by parenthesized text. Species without a specific name (ex: Nosema sp.) were numbered to give each entry a unique name.

To identify additional species, we searched the journal databases Scopus, Web Of Science and JSTOR for papers published between 1977 and April 2021. The search terms (“microsporidia” OR “microsporidium” OR “microsporida”) were used for all searches. Papers describing new microsporidia species or further describing existing species were used. Moreover, we looked at citing articles for papers describing new species, to find papers further describing those species. Non-English papers were translated into English using Google Translate. An additional 134 species were identified from a database of compiled by Kirk(49). Papers describing these additional species were found by searching in the aforementioned databases, or with Google. Microsporidia attributes for species described in journal articles were recorded as described above. For a subset of papers where the full article could not be accessed, information was recorded from the abstract.

### Analysis of microsporidia discovery date and geographical location

For each species, the earliest year from the “Date Identified” column of Table S1 was extracted. Species without identification year data were excluded from this analysis. Data of the countries where each species had been reported was extracted from the “Locality” column of the database. A species found in multiple countries was counted for all of those countries. State and province distribution maps for the United States and Canada were created using the same method. With the exception of oceans, species found in bodies of water were counted as the country located closest to it. If a species was reported either in a country that no longer exists or a general region, it was counted as the largest country that occupied its lands. For example, species found in the U.S.S.R. were counted as species found in Russia. Species reported from each continent were determined according to the categorization of worldpopulationreview.com.

### Determination of host taxonomy and environment

The taxonomy for each host species was determined using Catalogue of Life (http://www.catalogueoflife.org/annual-checklist/2019/), GBIF (https://www.gbif.org/), WoRMS (http://www.marinespecies.org/), NCBI Taxonomy Browser (https://www.ncbi.nlm.nih.gov/Taxonomy/Browser/wwwtax.cgi) or Wikipedia (https://en.wikipedia.org/). Using this approach, we annotated the taxonomy for 1422 out of 1449 host species. Catalogue of Life was used to determine the number of extant species in each phyla and the number of metazoan phyla, with the exception that Myxozoa was considered part of Cnidaria.

Host environment was determined using Catalogue of Life, GBIF and WoRMS. Host environment data was also extracted from other online databases, papers describing the host, or the original papers describing infection of the host by microsporidia (Table S2). To classify moths, all non-Pyralidae moths were classified as “terrestrial”. Pyralidae moths were classified as freshwater and terrestrial, since they have aquatic larval stages(65). If environment data was not available at the species level, we used environment data from the closest taxonomic level, the highest being order. Some species without environment data in the sources mentioned above, were assigned environments by common knowledge, i.e., parakeets inhabiting a terrestrial environment. The environment of 1338 out of 1449 species was annotated. Experimental hosts for microsporidia were excluded from analysis, as they may not reflect the natural host range of a microsporidia species(66).

### Categorizing and counting tissues infected

All unique infected-tissue names were extracted from the “Site” column of Table S1. These names were sorted into 23 broader tissue categories (Table S3A). Each unique tissue name was restricted to a single tissue category. Tissues that could not be clearly assigned to a category were classified as “Ambiguous”. Infections of protist species were classified as “Protist”. To count the number of tissues infected by each microsporidium, we counted how many tissues were listed for each microsporidium. Microsporidia species with systemic or ambiguous infections were excluded, as tissues in these categories could not be associated with a single tissue. The overall infection frequencies within tissue categories were determined using a custom R script. Each species was counted as infecting a tissue category one or zero times. Ambiguous and protist infections were not counted.

### Comparing infected hosts to infected tissues

To compare the number of hosts infected to the number of tissues infected, microsporidia species were grouped based on the number of hosts they infect. The proportion of species infecting one tissue was calculated within each host bin. Species with systemic infections were counted as infecting multiple tissues. Experimental hosts were excluded from analysis.

### Determining microsporidia transmission mode

Microsporidia’s mode of transmission was either determined from literature transmission data or inferred from the tissue infected, as has been done previously(43). (“Site” and “Transmission (described in literature)” columns in Table S1). Species without literature transmission data were classified as vertically transmitted if infecting germline, horizontally transmitted if infecting somatic tissue, or both. Transmission mode was not determined for species with systemic infections, as it is unclear what tissues they infect. When available, literature transmission data supplemented transmission mode determination. Species only infecting somatic tissues but with vertical transmission data from literature were classified as both. Species only infecting germline with horizontal transmission from literature were also classified as both. For species without tissue infection data, mode of transmission was determined solely through literature transmission data (horizontal, vertical, or both). Transmission mode was not determined for species with no tissue data and only autoinfection as literature transmission data, due to uncertainty over its mode of transmission. Moreover, transmission mode was not determined when transmission was explicitly described as ambiguous in literature. When comparing rates of horizontal and vertical transmission across environments, transmission modes were described for individual microsporidia host species. Only hosts in single environments were considered. Host species that could not be matched to specific tissues or modes of transmission were excluded from analysis.

### Spore shape classification

Spore shapes were classified using spore shapes reported in each publication and extracted from the “Spore Shape” column in Table S1. These shapes were grouped and placed into 20 shape categories (Table S3B). Shapes that could not be categorized as a defined grouped shape were labelled “ambiguous” and excluded from analysis. If a given spore was documented to be oval in addition to another shape, the latter shape classification was kept and used for analysis. However, if a spore was documented to be oval in addition to at least two other shapes, it was excluded from analysis.

### Analysis of spore dimensions

Spore lengths and widths were extracted from “Spore Length Average” and “Spore Width Average” columns in Table S1. If a given species had multiple measurements for a spore class, the “fresh” measurement was used and the remaining “fixed” measurements were excluded. If a spore class only had “fixed” measurements available, they were included in the analysis.

### Calculating spore volume

Spore volume was estimated for all types of documented spores using the formula for an ovoid, as this shape accounts for the majority of spores (Fig. 4A). The following formula was used: Volume= 4/3πabc, where a, b, and c are the 3 semi-axes of the ovoid. Width/2 was used for both a and b as width and depth of spores have been shown to be similar(41). Length/2 was used for c. When possible, priority was given to the “fresh” measurements of documented spore for calculating volume.

### Calculating polar tube length

Polar tube length was calculated by using the spore width and number of observed polar filament coils documented. If a range of coils was provided instead of a single value, the average number of coils was used. Spore width typically came from light microscopy observations, while coils came from transmission electron microscopy. Fresh width values were prioritized for calculations. The following formula was used: *Calculated polar tubule length = π* × width of spore × number of polar filament coils. Polar tube length was not calculated for spores with uncoiled polar tubes.

### Microsporidia tissue specificity

To assess if microsporidia polar tube length affects which tissues are infected, hosts infected by multiple microsporidia species were identified. Calculated polar tube lengths were organized into “Intestinal” infection if the species only infected the gut/digestive system as determined in Table S3A. If a species infecting the same host infected non-intestinal tissues, it was classified as “Other Tissues”. Every pair of microsporidia species between these two classes was compared for every host (Table S4).

### Comparing polar tube length to volume

To compare polar tube length to spore volume, only species with experimental polar tube values were considered. Species with multiple spore classes were represented multiple times. Entries where spore class could not be clearly matched to volume, polar tube length, or nucleus data were excluded. Least squares regression was performed for all spores (Fig. S4), as well as uninucleate and binucleate spores individually to control for the confounding effects of nuclei number. Correlation was calculated with Spearman r_s_ to limit the effect of outliers on correlation values.

### 18S phylogenetic analysis of microsporidia species

Using a modified version of the *read*.*Genbank* function from the *ape* package for R, microsporidia 18S sequences and associated data were collected from the NCBI GenBank nucleotide database: *https://www.ncbi.nlm.nih.gov/genbank/*. The search terms *(“Microsporidia”[Organism] AND (rRNA[Title] OR ribosomal[Title])) AND (small[Title] OR SSU[Title] OR 18s[Title] OR 16s[Title])* were used to collect 7830 unique 18S sequences representing 1107 unique microsporidia species identifiers. The longest sequence for each unique species identifier was extracted from this collection for subsequent analysis. Only the sequences with species identifiers corresponding to species in our database (Table S1) were included in subsequent steps. This left us with 18S sequences for 273 unique microsporidia species. We also included a single 18S sequence for *Rozella allomycis*, a commonly used outgroup in microsporidia phylogenies(67). A list of accession numbers for all 274 18S sequences and corresponding species can be found in Table S5A.

The 274 18S sequences were aligned with MAFFT, using the default parameters. The alignment was masked using the *MaskAlignment* function from the *DECIPHER* package for R. Bayesian phylogenetic analysis of the masked alignment was performed with MrBayes v 3.2.7, using two MCMC runs, each with one cold and three heated chains, for 10 million generations, sampling every 1000 generations, with the first 2.5 million generations being discarded as burn-in. For this inference, we used the GTR model of substitution, with an *“invgamma”* model for rate variation. Maximum Likelihood phylogenetic analysis of the masked alignment was performed with RAxML-HPC2 using the default parameters and 1000 bootstrap iterations. 50% majority rule consensus trees were generated for both Bayesian and Maximum Likelihood analyses and were annotated using the *ggtree* and *treeio* packages for R. Clade groupings from both trees were compared by computing the fraction of species placed in the same clade in both trees. A complete list of parameters for the MrBayes and RAxML phylogenetic analyses can be found in Table S5B. MAFFT, MrBayes and RAxML were all run on XSEDE accessed via the CIPRES Science Gateway: *https://www.phylo.org/*(68).

### Analysis of microsporidia evolutionary relationships

All analysis of evolutionary relationships between the four major clades was done in R. The distribution of host environments was determined by counting the number of species in each clade that are found in each environment, expressed as a percentage of the total number of species in each clade. The distributions of spore volume and polar tube length were determined using the calculated spore volume and calculated polar tube length data (Table S1). In cases where species had multiple spore classes, all were included as separate data points in the analysis. To determine if any hosts were infected by multiple microsporidia species from different clades, the assortment of hosts for each clade were compared with each other to see if any hosts appeared in multiple clades.

Using the *dist.alignment* function from the *seqinr* package for R, a pairwise sequence similarity matrix for the 36,315 unique species pairs encompassing 270 microsporidia species from the phylogenetic analysis was computed based on the unmasked MAFFT alignment. (The unnamed Pleistophora sp. 1, 2 and 3 were excluded due to ambiguity in species identity between our database and GenBank). The fractions of species pairs within a given sequence similarity range that had at least one shared host taxonym, environment, and infected tissue were computed separately and displayed as distributions with 20 sequence similarity bins. The percentage similarities in spore volume and polar tube length in each species pair were computed separately according to the formula *(larger value / smaller value)*100*, using the calculated spore volume and calculated polar tube length data (Table S1). In cases where species had multiple spore classes, values for each class were averaged prior to the percentage similarity computation. The average percentage similarities in spore volume as well as polar tube length within a given sequence similarity range were then computed and displayed as distributions with 20 sequence similarity bins. For each distribution, species pairs that had at least one species with missing data for the given comparison criterion were excluded.

## Data and code availability

All microsporidia species and host data are available in Tables S1 and S2. The most recent version of the microsporidia species data can be at www.placeholder.com. All data analysis was performed using Microsoft Excel and R version 3.6.1 or later accessed via RStudio version 1.2.5019 or later(69). All code used for data analysis can be found on the Microsporidia-Database GitHub repository *https://github.com/bmurareanu/Microsporidia-Database*.

## Acknowledgements

We thank Hala Tamim El Jarkass, Alexandra R. Willis, JiHae Jeon, and Robert J. Luallen for providing helpful comments on the manuscript. We thank Alexandra R. Willis for providing the illustrations for figure 2. This work was supported by the Natural Sciences and Engineering Research Council of Canada (Grant #522691522691 and Undergraduate Student Research Awards to B.M., R.S., R.Q., and J.J.) and an Alfred P. Sloan Research Fellowship FG2019-12040 (to A.W.R.).

## Author contributions

B.M., R.S., R.Q., J.J, and A. R. designed the project, planned the data analysis, and co-wrote the paper. B.M., R.S., R.Q., and J.J. carried out all of the data curation and analysis.

## Supplemental material

**Fig. S1. Orders of Arthropods infected by microsporidia**.

**Fig. S2. Relationship between microsporidia spore dimensions and shape classification**.

**Fig. S3. Relationship between polar tube length and spore volume**.

**Fig. S4. Microsporidia host environment and spore morphology are independent of infection transmission mode**.

**Fig. S5. Bayesian 18S phylogeny of microsporidia species**.

**Table S1. Database of microsporidia species attributes**.

**Table S2. Taxonomical descriptions of microsporidia hosts**.

**Table S3. Microsporidia host-tissues and spore-shape classifications**.

**Table S4. Polar tube lengths of species pairs that infect only the intestine and also other tissues in a common host**.

**Table S5. Accession numbers for microsporidia DNA sequences and parameters for MrBayes phylogenetic analysis**.

**Fig. S1.**
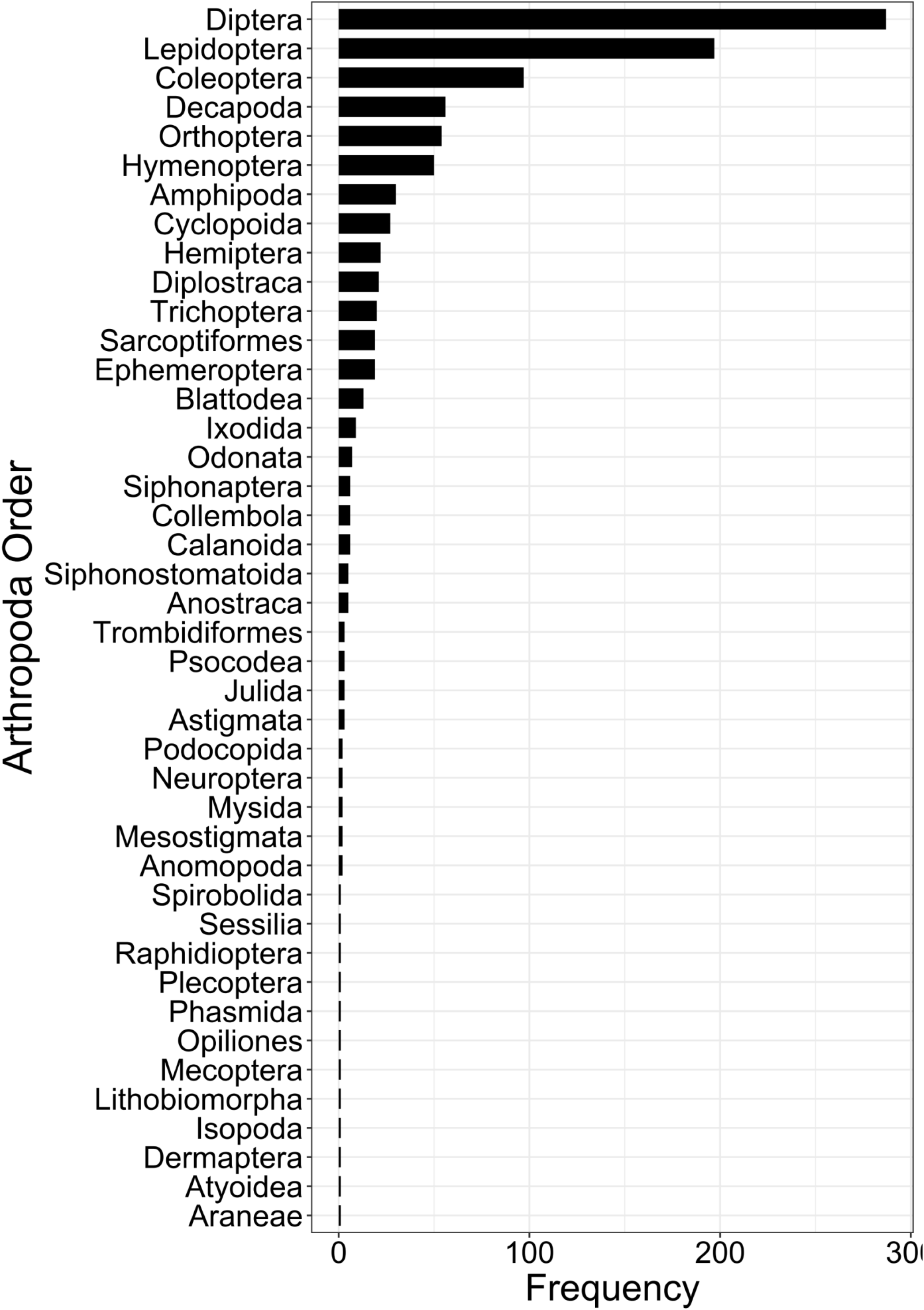
Frequency of Arthropod order infections. Frequency of Arthropod orders reported to be infected by microsporidia (n = 46 orders).

**Fig. S2.**
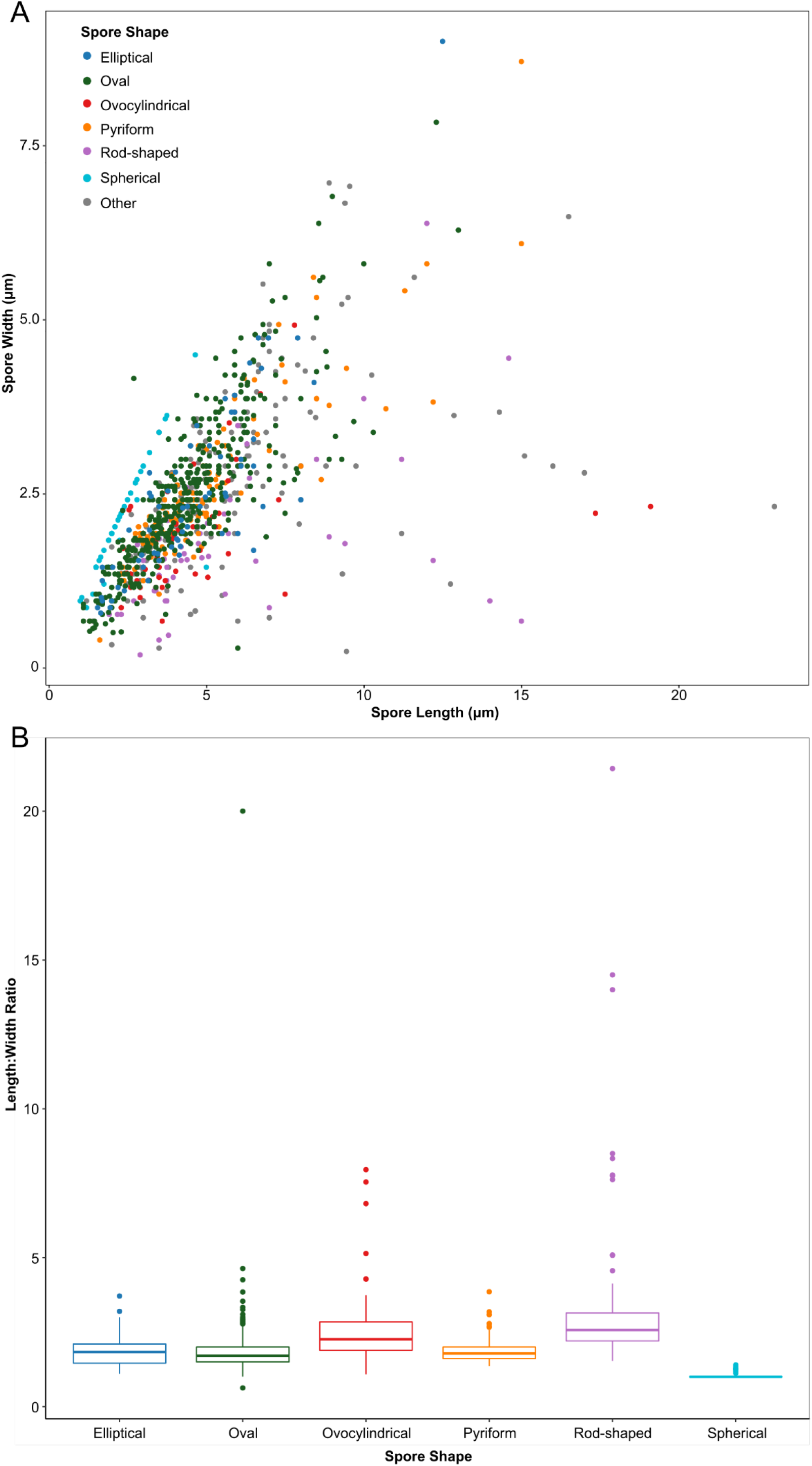
Relationship between microsporidia spore dimensions and shape classification. **(A)** Spore length and width dimensions for the six most common shape groups (Table S4) with the 16 remaining shapes classified as other (n=1032 spores). **(B)** Spore length to width ratios for the six most common shape groups.

**Fig. S3.**
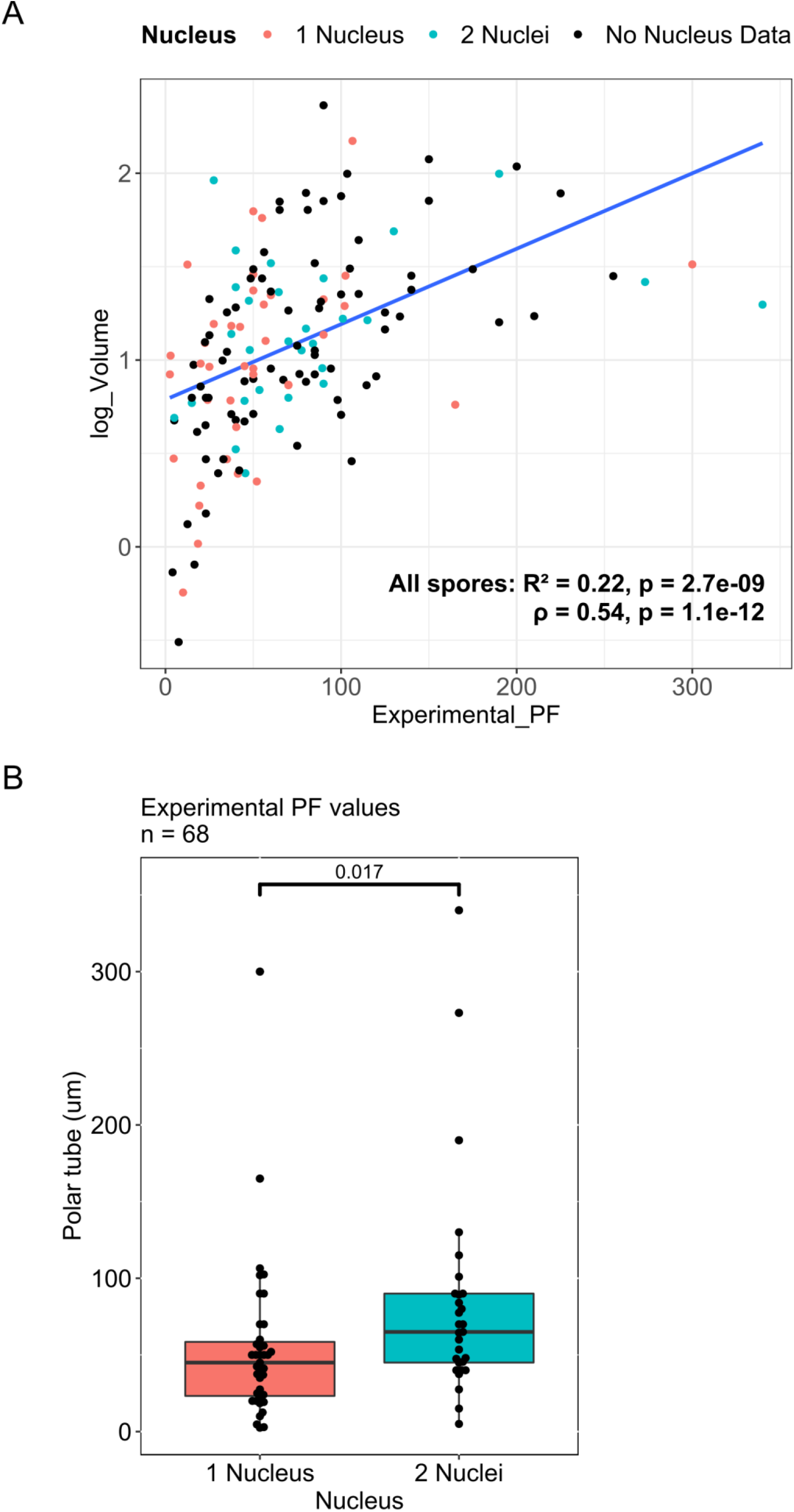
Relationship between polar tube length, nuclei number and spore volume. **(A)** Comparison between experimental polar filament lengths and spore volume for spores of any nuclei number (n = 147). Correlation was calculated with spearman r_s_. **(B)** Median spore experimental tube length for uninucleate (n = 39) and binucleate (n = 29) spores. The p-value was determined by the Mann–Whitney U test.

**Fig. S4.**
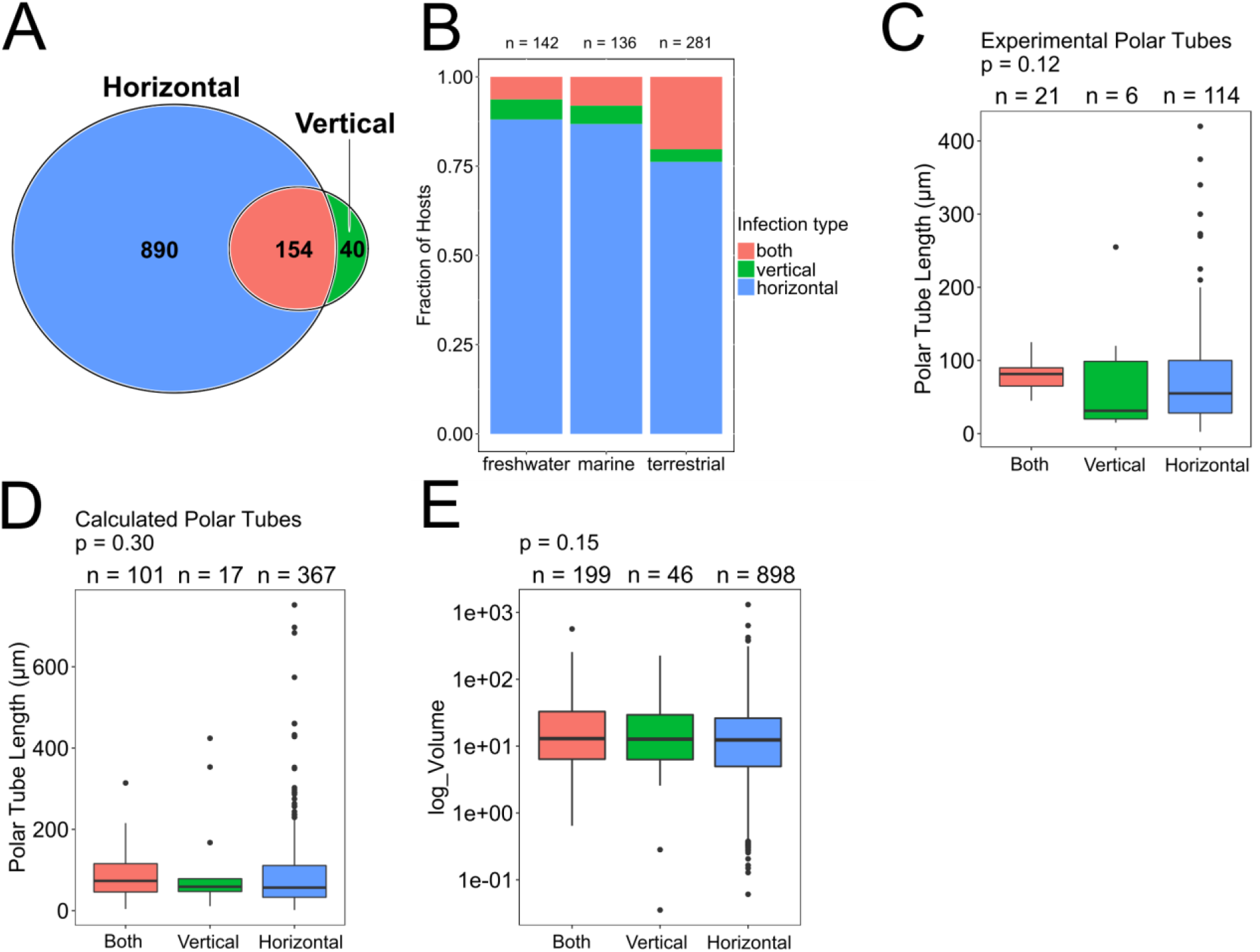
Microsporidia host environment and spore morphology are independent of infection transmission mode. The transmission mode of each species of microsporidia was inferred from tissues infected, with those that only infect somatic tissues being classified as horizontally transmitted, those that only infect germline tissues being classified as vertically transmitted, and those that infect both types of tissues being classified as both (see methods). **(A)** Venn diagram of transmission mode of microsporidia species (n = 1084 microsporidia species). **(B)** Comparison of infection transmission mode to host environment (n = 559 hosts). Only hosts classified as belonging to a single environment were analyzed. **(C)** Comparison of infection transmission mode to experimentally measured polar tube values (n = 141 spores) and **(D)** calculated polar tube values (n = 485 spores). Species with multiple polar tube values were represented multiple times. **(E)** Comparison of infection transmission mode to spore volume (n = 1143 spores). Species with multiple spore sizes were represented multiple times.

**Fig. S5.**
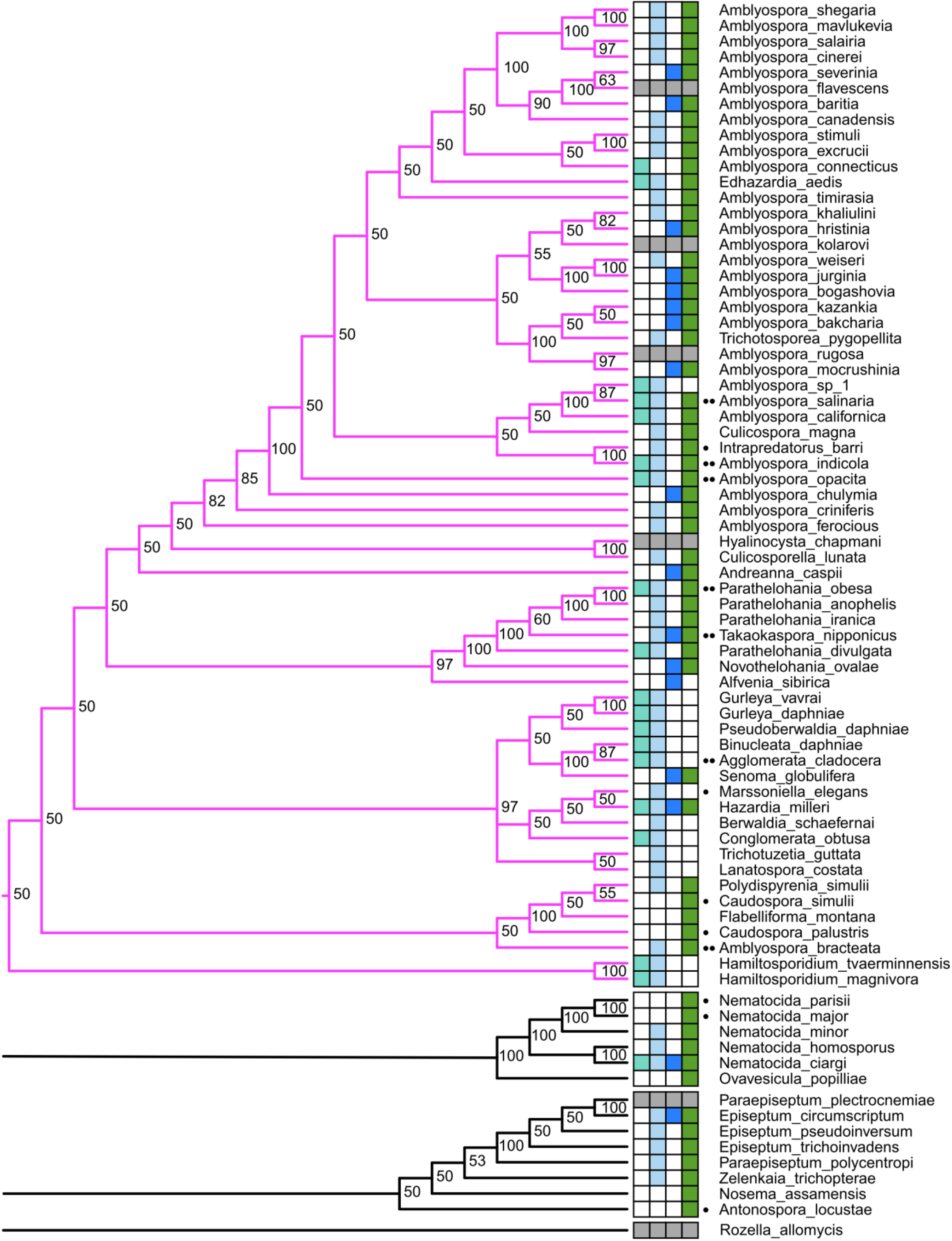
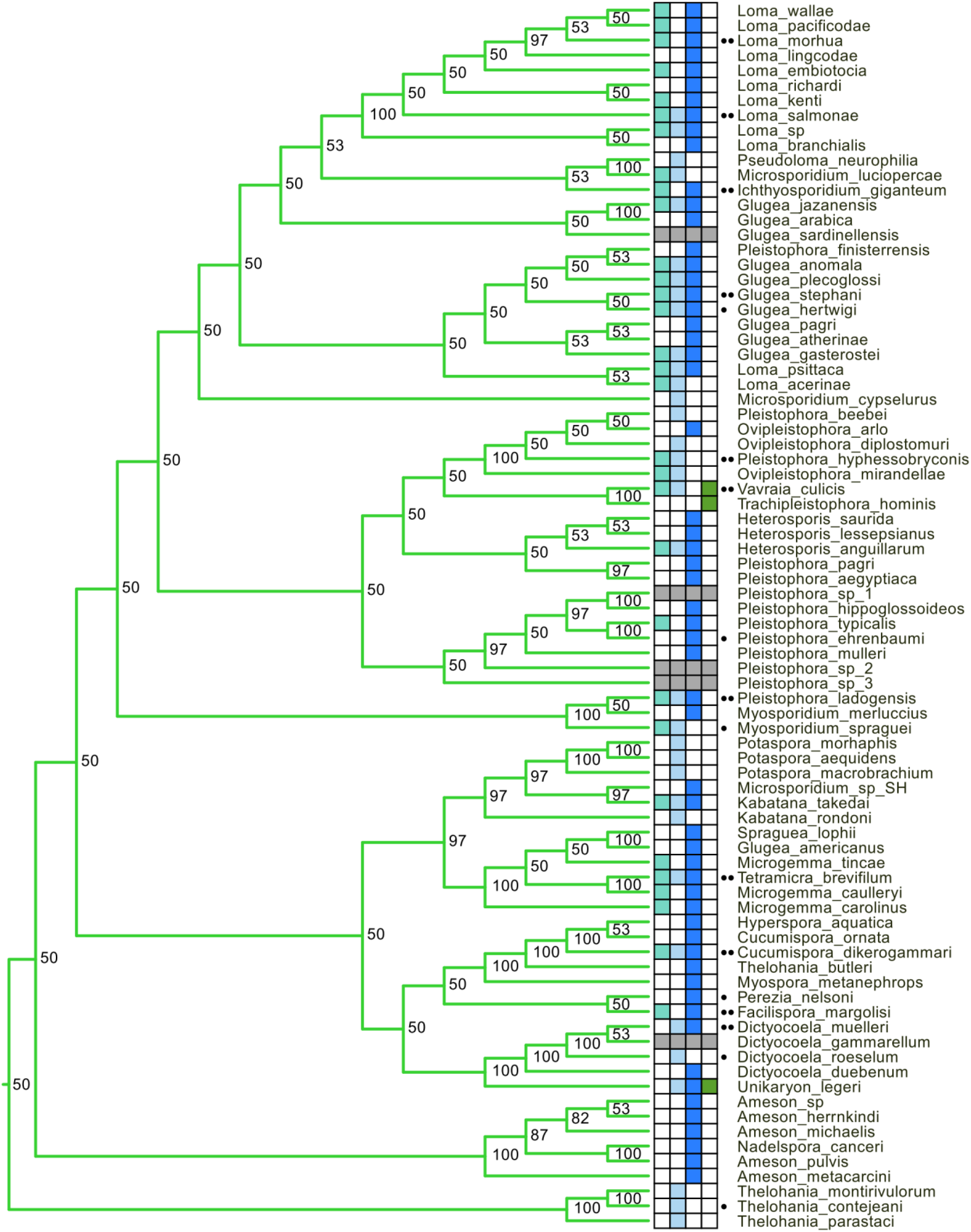
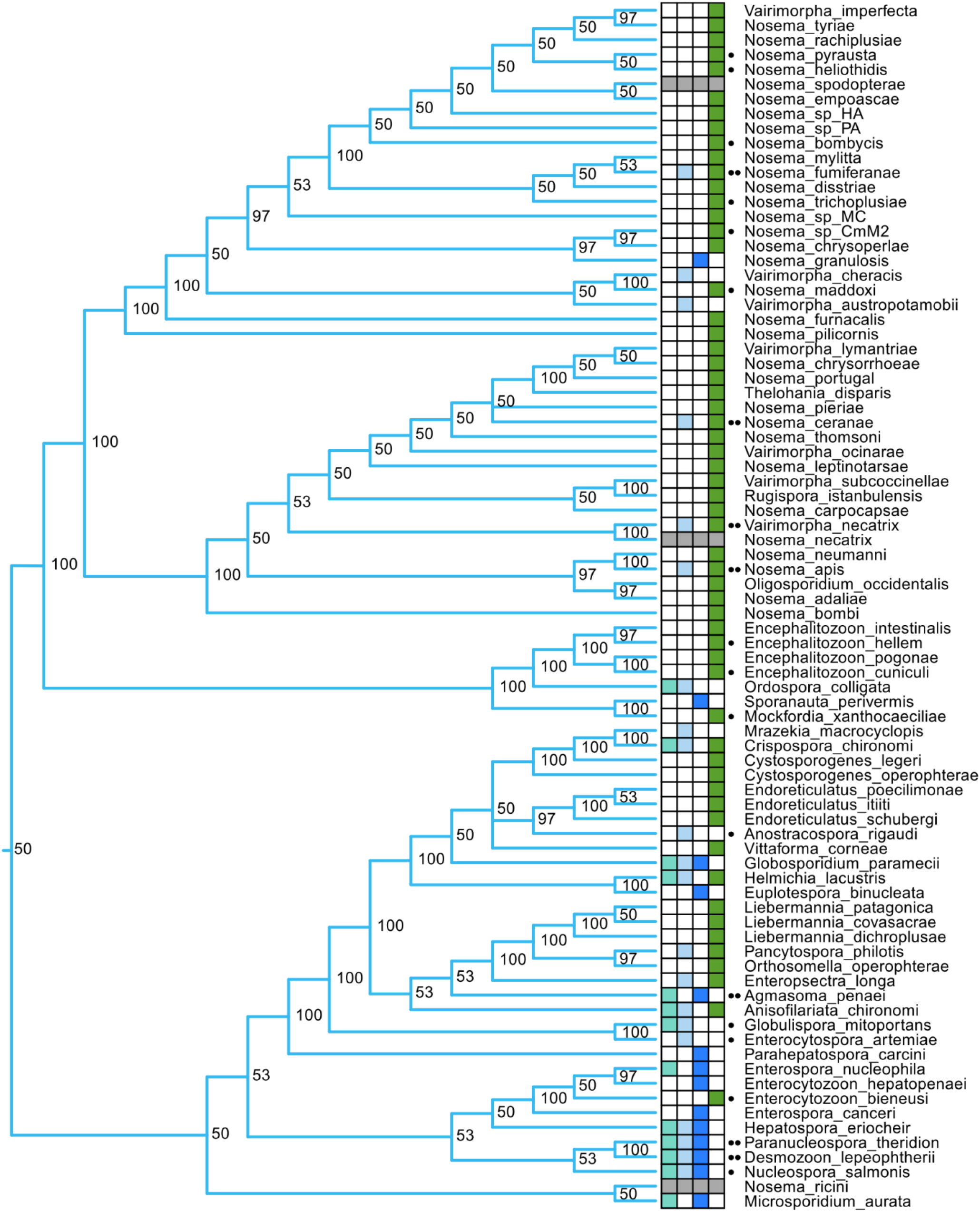
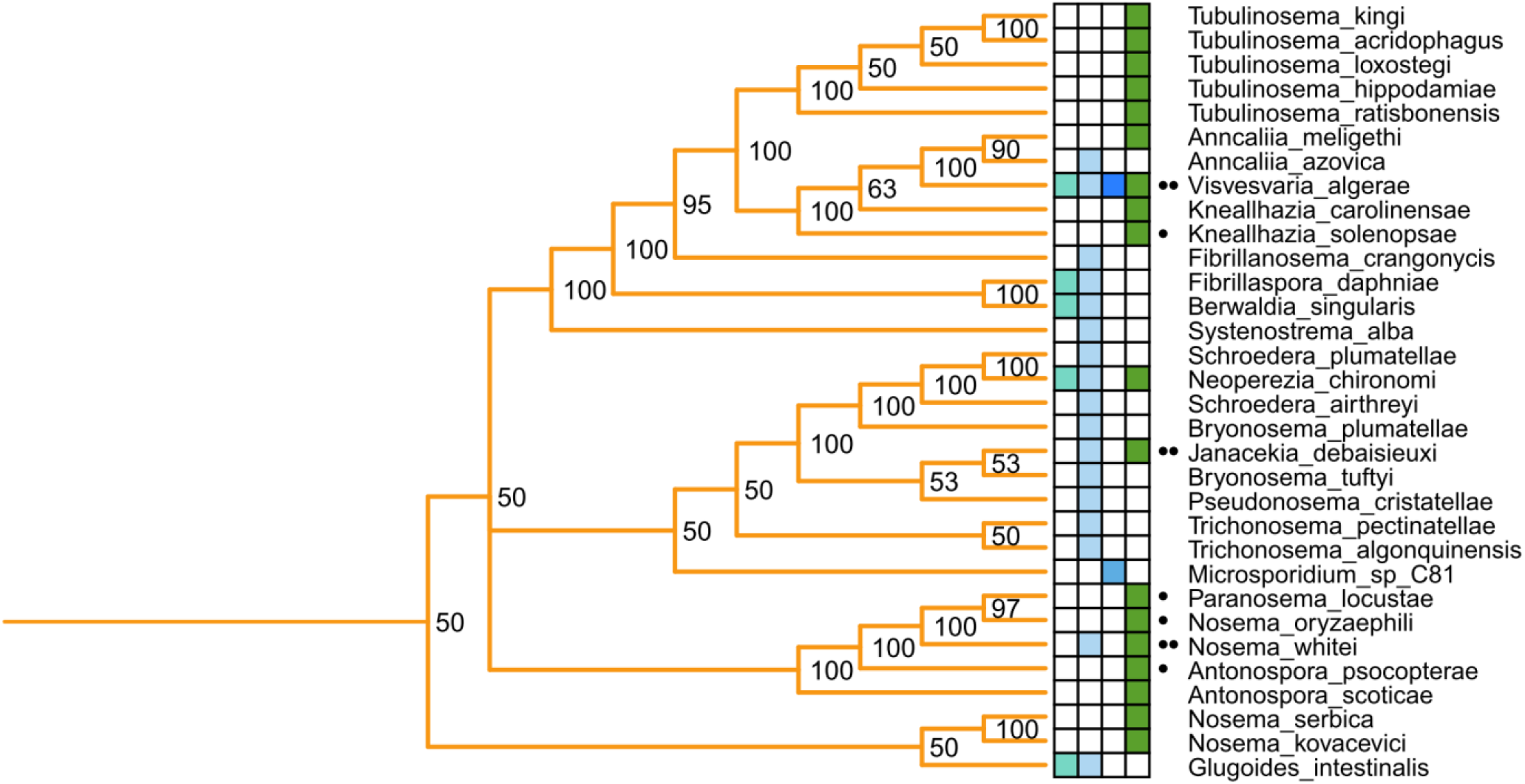
Bayesian 18S phylogeny of microsporidia species. Phylogenetic tree of 273 microsporidia species and *Rozella allomycis*. The tree was generated and labelled for clades and environments the same as in Fig 5. The species name of each branch is shown on the right, and the Bayesian posterior probability support value is shown at each node.

